# Ancestry-specific rewiring of BCR-MAPK signaling in sarcoidosis B cells

**DOI:** 10.64898/2026.04.20.718985

**Authors:** Christopher Dunn, Caleb Watkins, Gideon Hallum, Nathan Pezant, Astrid Rasmussen, Patrick M. Gaffney, Harini Bagavant, Umesh Deshmukh, Courtney G. Montgomery

## Abstract

Sarcoidosis is a heterogenous disease of unknown etiology characterized by non-caseating granulomas. Disease prevalence and presentation vary significantly by ancestry and ranges from acute, self-resolving disease to severe, chronic disease. Following previous reports suggesting B cells in the development and pathogenesis of sarcoidosis, we present here results of single-cell RNA sequencing, supporting B cell involvement in sarcoidosis through altered immediate early response, rewiring of MAPK signaling, and ancestry-specific preferential expansion of B cell receptors. Peripheral blood mononuclear cells were obtained from individuals of African or European Ancestry (AA and EA, respectively) including 48 healthy controls, 59 sarcoidosis patients, and 28 systemic lupus erythematosus (SLE) patients. SLE samples were used as a disease control. Differential expression analysis highlighted many differentially expressed genes (DEGs) with almost 5x more in the AA sarcoidosis versus AA control group compared to the EA sarcoidosis versus EA control group. B cells had the most DEGs of all cell types and expression patterns were similar between ancestries, however, sarcoidosis had an opposite transcription pattern than SLE, demonstrating an alternative immune response to acute activation than that seen in a prototypical autoinflammatory disease. This trend was maintained when examining specialized B cell subsets, with the most pronounced effect in the AA sarcoidosis versus AA control comparison. Our results strongly support further investigation of the role of humoral immune response in sarcoidosis and the potential to highlight patient groups likely to benefit from existing B cell therapies.

---

Sarcoidosis is a multi-system granulomatous disease marked by variable presentation and disease course. It occurs worldwide and affects both sexes^1^, with higher incidence and severity in Americans of African ancestry (AA) compared to Americans of European ancestry (EA)^2^. Sarcoidosis is characterized by a dysfunctional immune response and while literature has largely focused on T cells and monocytes/macrophages, an important role of B cells in sarcoidosis has long been suggested, including hypergammaglobulinemia, circulating immune complexes, autoantibodies, and sarcoidosis-lymphoma syndrome^3, 4^. Consistent with this, dysregulation of intercellular signaling pathways that regulate immune activation, including the MAPK pathway, has similarly been implicated in sarcoidosis^5^. However, these studies have stopped short of identifying a unified mechanism linking B cell abnormalities, MAPK signaling, and sarcoidosis.

In a recent interrogation of circulating immune cells in AA and EA sarcoidosis patients via single-cell RNA (scRNA) sequencing, we identified a signal-adapted transcriptional profile in B cells. This profile, including evidence of attenuated immediate-early response and MAPK feedback rewiring has previously been described in cancer-adjacent immune states and is present in both ancestries; however, it is much more pronounced in AA sarcoidosis, an effect likely to be missed in a heterogeneous sarcoidosis cohort. Additionally, we observed preferentially-expanded BCR families that appear unique to AA sarcoidosis further suggesting an ancestry-specific response in sarcoidosis B cells.

## Methods

Our single-cell dataset comprised 48 healthy controls (31 AA/17 EA), 59 sarcoidosis cases (15 AA/44 EA), and 28 systemic lupus erythematosus (SLE) patients (28 AA). Evaluation and consent of sarcoidosis patients and healthy controls were described previously, but, in brief, classification required biopsy proven sarcoidosis and the exclusion of autoimmune conditions, tuberculosis, or cancer^6^. SLE participants were enrolled to the Lupus Family Registry and Repository, managed by the Oklahoma Rheumatic Disease Research Cores Center^7^ from which PBMCs and partnered clinical data were obtained. scRNA sequencing was performed as previously described^6^, albeit with most recent versions of software.

The data was filtered (*Seurat*) including data-driven adjustments; cells with <500 features (genes), <1000 read counts, >12% mitochondrial genes, and >5% ribosomal genes were removed, as were estimated doublets (*Scrublet*). To eliminate red blood cell contamination, cells with >0.25% expression of HBB, HBA1, and HBA2 were removed. Cell clustering was performed on a dataset of 582,213 cells from sarcoidosis cases (248,963 cells), healthy controls (209,451 cells), and SLE cases (88,640 cells) (*Seurat*). Annotation was carried out with the MonacoImmuneData (*celldex*) dataset, comprising 114 human RNA-seq samples of sorted immune cell populations. Immune cells were first classified into 7 broad cell types (Monocytes, Dendritic cells, NK cells, T cells, CD8+ T cells, CD4+ T cells, and B cells) (**Figure 1**). These groups were then classified into specialized subsets resulting in 18 subgroups.

**Figure 1.**
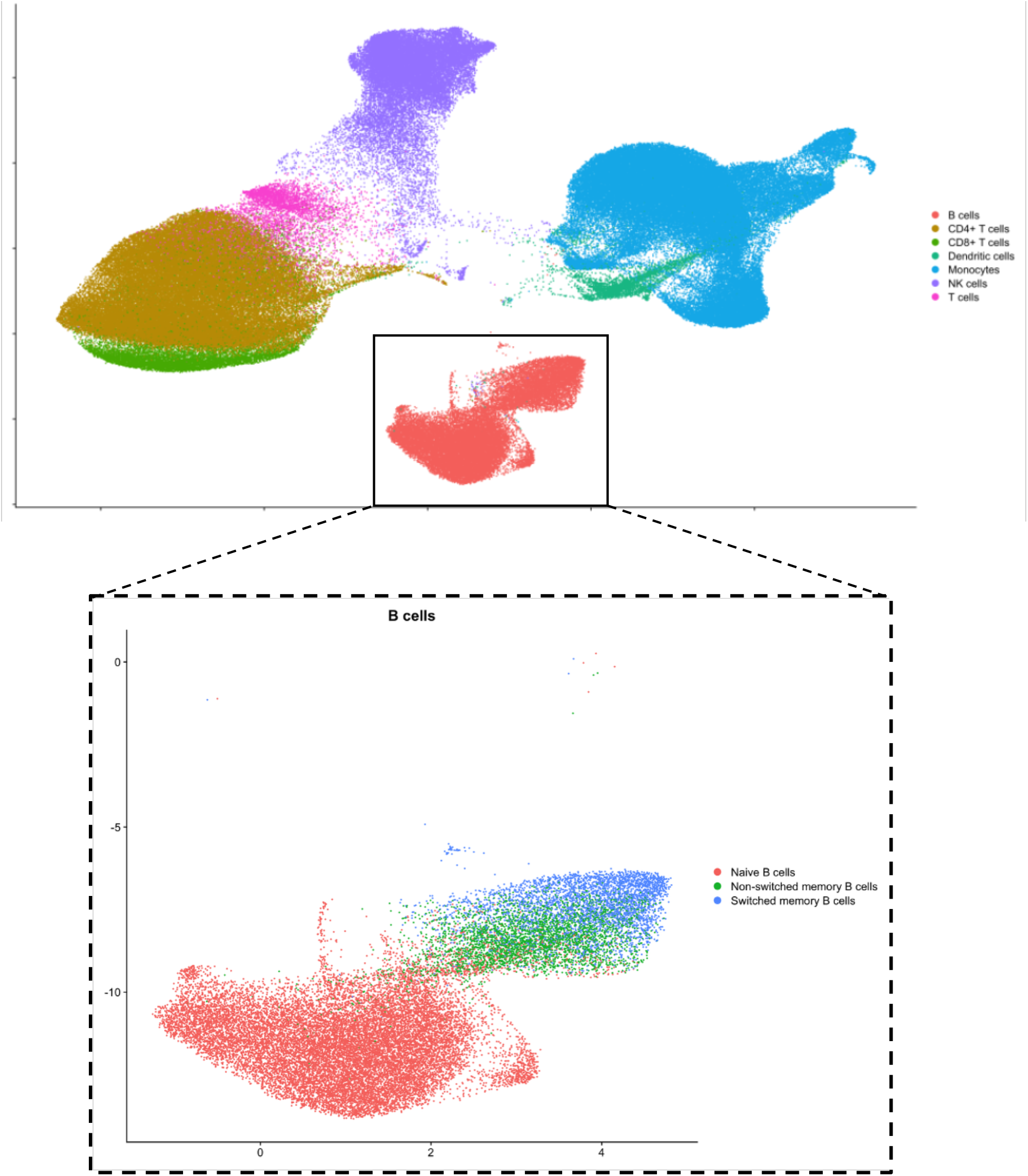
UMAP of immune cell clusters

To balance cell counts, a subset of each of the three participant groups was chosen for differential expression (DE) analysis and matched for age and sex. Association tests were performed with gender as a covariate, within ancestry, using logistic regression at a cell-specific level (*FindMarkers, Seurat)*. Genes with log2FC≥|1| and P_adj_ < 0.05 were considered significantly DE.

## Results and Discussion

DE analyses yielded significantly DE genes in all cell types, and in both ancestries. We found 5-fold more DE genes in AA than EA. Specifically, B cells had the most DE genes, both up- and downregulated, and due to the current lack of understanding of B cells in sarcoidosis we investigated the transcriptional pattern of these cells. As a point of comparison, we performed DE analysis between AA SLE and the same AA controls. SLE was chosen because it has a well characterized inflammatory profile. As expected, we found an opposite transcriptional pattern in AA SLE, suggesting transcriptionally attenuated B cell states in sarcoidosis rather than acute activation as seen in SLE (**Figure 2**). To better characterize this, we investigated DE results from our specialized B cell subsets including naïve B cells, non-switched memory B cells, and switched memory B cells. DE genes converged on three tightly linked biological themes: 1) attenuated immediate-early response, 2) altered MAPK/ERK feedback control, 3) and preferentially-expanded BCR genes (**Figure 2**).

**Figure 2.**
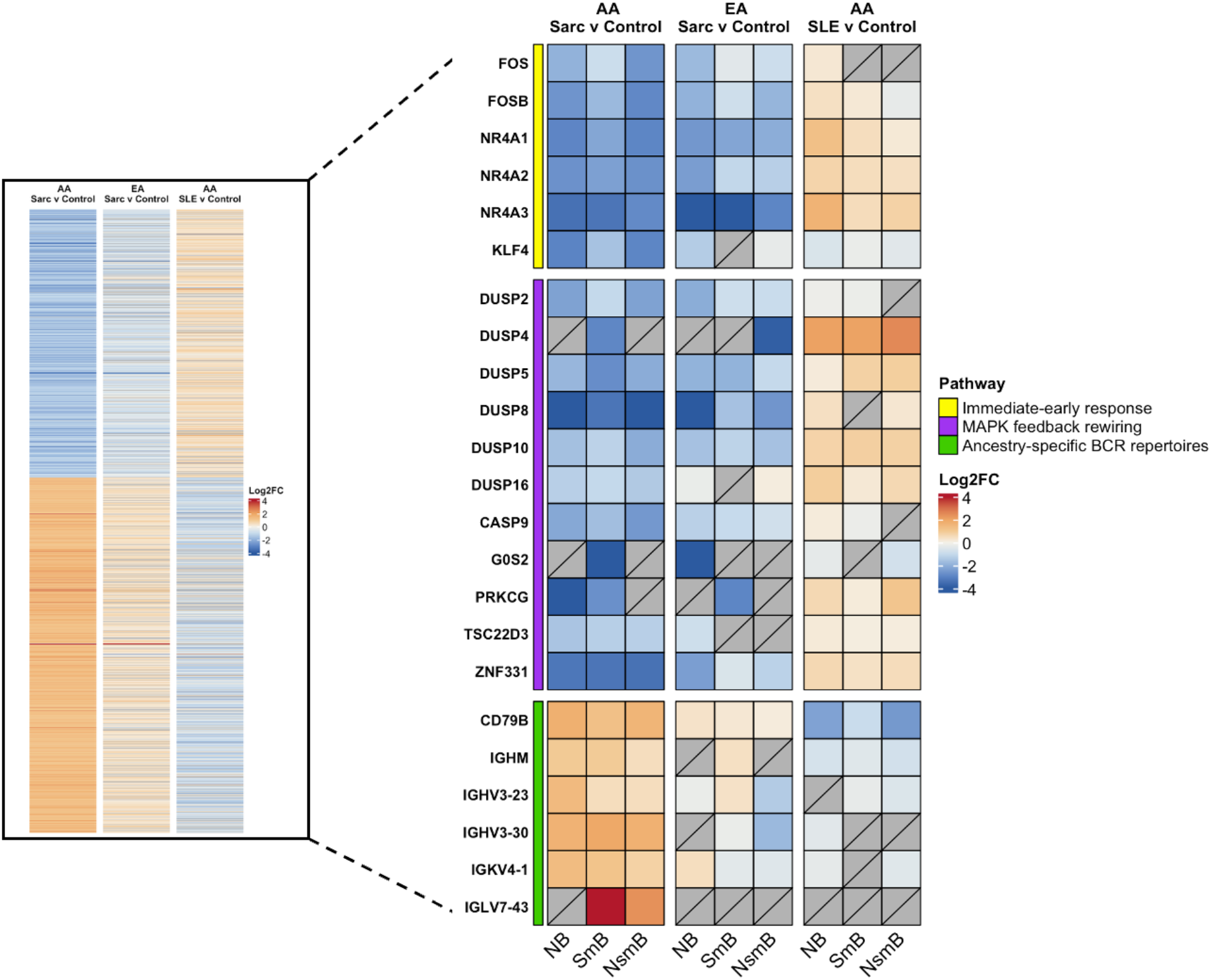
Heatmap of all statistically significantly differentially expressed genes from all B cells in our three comparison groups (left) and heatmap of genes specifically demonstrating B cell activity within B cell subsets (right).

First, we see coordinated loss of major transcription factors involved in immediate-early response including the FOS and NR4A family of genes as well as the stress-response transcription factor KLF4. Second, although MAPK/ERK has been implicated in sarcoidosis^8^, our observation of ERK negative feedback regulation failure suggests a deviation from typical hyperactivation seen in other immune-mediated diseases, such as SLE. Specifically, downregulation of multiple dual-specificity phosphatase genes (DUSP2, DUSP4, DUSP5, and DUSP8, DUSP10, DUSP16), which normally serve as ERK-inducible negative feedback regulators and as tumor suppressors. Loss of these regulators alters ERK signaling, leading to dysregulated, sometimes sustained, signaling states^9^. Within this group of genes, we see some similarity in expression between AA and EA, suggesting a shared pathological signaling mechanism; however, the pattern is much more pronounced in AA patients, implicating ancestry-specific genetic or regulatory differences in stress response, MAPK feedback, and B cell survival. Finally, we see coordinated upregulation of several BCR genes including CD79B, and to a lesser extent CD79A (n.s.), that are associated with heightened BCR signaling. IgM upregulation in sarcoidosis for both ancestries suggests a bias as the other immunoglobulin genes were not significantly DE. We also see evidence of restricted immunoglobulin usage with upregulation of IGHV3-23, IGHV3-30, IGKV4-1, and IGLV7-43.

Our results are the first to offer a link between B cell abnormalities, MAPK signaling, and sarcoidosis. They include the first report of an ancestry-specific expansion of BCR genes in sarcoidosis and highlight the potential of immunologically informed treatment regimens, including the use of B cell therapies shown to be effective in refractory disease^10^. They further demonstrate the potential in using indicators of humoral immunity as biomarkers for disease and a means by which to stratify patients to elucidate disease mechanisms. Finally, our study points to the continued need for molecular characterization of sarcoidosis in patients across genetic backgrounds and disease course.

## Acknowledgments

Christopher Bottoms, Cherilyn Pritchett, Judy Harris, and participants from cohorts

